# Improving Multichannel Raw Electroencephalography-based Diagnosis of Major Depressive Disorder by Pretraining Deep Learning Models with Single Channel Sleep Stage Data^*^

**DOI:** 10.1101/2023.05.29.542700

**Authors:** Charles A. Ellis, Abhinav Sattiraju, Robyn L. Miller, Vince D. Calhoun

## Abstract

As the field of deep learning has grown in recent years, its application to the domain of raw resting-state electroencephalography (EEG) has also increased. Relative to traditional machine learning methods or deep learning methods applied to extracted features, there are fewer methods for developing deep learning models on small raw EEG datasets. One potential approach for enhancing deep learning performance in this case is the use of transfer learning. In this study, we propose a novel EEG transfer learning approach wherein we first train a model on a large publicly available sleep stage classification dataset. We then use the learned representations to develop a classifier for automated major depressive disorder diagnosis with raw multichannel EEG. We find that our approach improves model performance, and we further examine how transfer learning affected the representations learned by the model through a pair of explainability analyses. Our proposed approach represents a significant step forward for the domain raw resting-state EEG classification. Furthermore, it has the potential to expand the use of deep learning methods across more raw EEG datasets and lead to the development of more reliable EEG classifiers.

**Clinical Relevance:** The proposed approach takes the field of deep learning in EEG a step closer to the robustness needed for clinical implementation.

## I. Introduction

The application of deep learning methods to raw resting-state electroencephalogram (EEG) data has grown increasingly common in recent years. However, given the relatively small size of many locally collected EEG datasets and the expense of collecting larger datasets, it can be challenging to train reliable EEG deep learning models. It would be ideal if large existing EEG datasets could be leveraged to improve performance in other EEG deep learning applications. Other areas of deep learning (e.g., image classification) often use transfer learning for that reason. Additionally, some EEG studies involving manually extracted features (e.g., spectrograms) have developed transfer learning approaches. Nevertheless, while some initial efforts have been made to develop pretraining methods for EEG, there remains significant opportunities for growth in that arena. In this study, we implement a novel approach to demonstrate how large publicly available raw single-channel EEG sleep stage classification datasets can be used to pretrain a convolutional neural network (CNN) for automated multichannel EEG-based major depressive disorder (MDD) diagnosis. We show that the representations learned by the pretrained CNN can be used to obtain well-above chance-level performance without any further tuning of the convolutional layers, and we further show how our pretrained model can yield higher performance when fully tuned relative to the same architecture without pretraining. We lastly apply explainability analyses to examine how pretraining affected the learning of representations for canonical frequency bands and channels.

Extracted features to raw EEG. Historically, a large number of studies have applied machine learning and deep learning methods to extracted EEG features [1], [2]. While these approaches can be effective, their use of manually extracted feature inherently limits the size of the feature space from which they can learn. As such, the application of deep learning methods to raw EEG has become increasingly common, as deep learning methods like CNNs have the capability to perform automated feature extraction [3], [4].

Unfortunately, relative to traditional machine learning models trained on manually extracted features, deep learning models require larger datasets to be effective. Large datasets can be very time- and resource-intensive to collect and may be outside the capabilities of smaller research centers. Within the space of raw EEG analysis, some studies have addressed this problem via the use of data augmentation with additive noise and segmentation with large overlaps between samples [5][6]. Another possible solution is the use of transfer learning.

Transfer learning involves the training of a model on a large dataset from one task. The representations learned from that dataset can then be reused in the initialization of model weights for a second task. Transfer learning is common in applications like image classification and has been used in multiple studies involving extracted EEG features [7], [8]. Nevertheless, there are relatively few pretraining approaches for raw EEG. Of those that have been developed, they often involve event-related potentials [9] or task data [10] that are very different from resting state data or the use of self-supervised learning [11]. One study of particular interest used a pretrained architecture originally developed in the imaging domain to classify raw EEG data [12].

In this study, we develop an approach for pretraining models on publicly available raw one-channel sleep stage classification data and subsequently applying those models for the classification of multichannel EEG data. We demonstrate our approach within the context of automated major depressive disorder diagnosis. We find that our approach is able to learn generalizable features and that our pretraining approach improves model performance relative to a baseline model with the same architecture and a traditional training approach. Lastly, we apply explainability approaches to examine differences in the features learned by each of our models.

## II Methods

### A. Datasets

We used two publicly available datasets in this study: a single-channel EEG sleep stage classification dataset and a multichannel EEG MDD dataset. Given that both datasets were publicly available, it was unnecessary to obtain approval for this study from an institutional review board.

#### Sleep Stage Dataset

We used a portion of the Sleep Cassette data from the Sleep-EDF Expanded dataset [13] on PhysioNet [14]. The dataset was composed of 153 20-hour recordings from 78 study participants, though due to computational constraints in the training process we used data from only 39 participants. The data was recorded at a sampling rate of 100 Hertz. While 2 electrodes were collected, we used data from the Fpz-Cz electrode. Thirty-second segments of data were assigned to Awake, Rapid Eye Movement (REM), Non-REM 1 (NREM1), NEM2, NREM 3, and NREM4 sleep stages by an expert.

Based on the expert annotations, we separated the data into 30-second segments. To alleviate class imbalances, we removed the Awake data at the start of each recording and many Awake segments at the end of each recording. According to existing clinical practice [15], we reassigned NREM4 to NREM3. We lastly z-scored samples from each recording. Our final dataset was composed of 37,605, 7,534, 35,262, 7,939, and 13,801 Awake, NREM1, NREM 2, NREM3, and REM samples, respectively.

#### MDD Dataset

We used a scalp EEG dataset [16] composed of 5-10 minute resting-state recordings with eyes closed from 28 healthy controls (HCs) and 30 individuals with MDD (MDDs). Recordings were performed at a sampling frequency of 256 Hertz in the standard 10-20 format with 64 electrodes. Similar to previous MDD studies, we used 19 channels: Fp1, Fp2, F7, F3, Fz, F4, F8, T3, C3, Cz, C4, T4, T5, P3, Pz, P4, T6, O1, and O2. We performed several data normalization steps to make the MDD data similar to the sleep data that would be used for pretraining. Specifically, we downsampled the data to 100 Hz and channel-wise z-scored each recording separately. To increase the size of the dataset available for training, we used a 30-second sliding window with a 2.5-second step size to epoch the data. Our final dataset contained 2840 HCs and 2850 MDDs.

### B. Model Development

We adapted the architecture (shown in Figure 1) that was used for MDD classification in [6], [17] and originally for schizophrenia classification in [3]. We trained 4 models using highly similar architectures: a basic MDD model with no pretraining (Model A), a sleep stage model (Model B) that would later be used for pretraining MDD models, a pretrained MDD model with frozen feature extraction layers (Model C) to examine the transferability of the features learned by the sleep model, and a pretrained model (Model D) that extended Model C by using the trained dense layer weights and training the whole network. MDD models A, C, and D used sensitivity (SENS), specificity (SPEC), and balanced accuracy (BACC) when assessing model performance, and the sleep model (Model C) used SENS, precision (PREC), and the F1 score.

**Figure 1.**
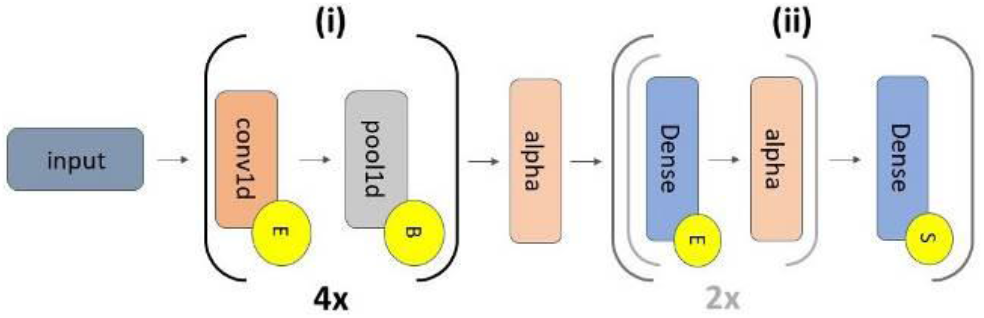
Model Architecture. Models A through D have 2 sections that are separated by an alpha dropout layer (alpha): (i) feature extraction, which repeats 4 times, and (ii) classification. The grey inset within section (ii) repeats twice. The 4 convolutional (conv1d) layers have 5, 10, 10, and 15 filters, respectively (kernel size = 10, 10, 10, 5). They are followed by max pooling layers (pool size = 2, stride = 2). Section (ii) has 3 dense layers (64, 32, and 2 nodes) with interspersed alpha dropout layers (alpha) with rates of 0.5. The alpha layer between (i) and (ii) also has a rate of 0.5. Yellow circles containing an “ E”, “B”, or “ S” correspond to ELU activations, batch normalization, and softmax activations. All conv1d and dense layers had max norm kernel constraints with a max value of 1. In Model B the channel dropout layer was placed between “ Input” and Section (i).

#### MDD Basic Model (Model A)

We used 10-fold stratified group shuffle split cross-validation to prevent samples from the same participant being distributed across training, validation, and test sets. Training, validation, and test sets were assigned approximately 75%, 15%, and 10% of the data in each fold, respectively. To increase dataset size, we applied a data augmentation approach that involved creating a copy of the training data in each fold, adding Gaussian (µ=0, σ=0.7) noise to the copy, and using the combined original and augmented data for training. This approach has been used in multiple previous studies [5]. We next used a class-weighted loss function to account for training set class imbalances. We applied the Adam optimizer (learning rate = 0.0075), a batch size of 128, 35 epochs, and early stopping if 10 epochs passed without an increase in validation accuracy.

#### Sleep Stage Model with Modifications for Pretraining (Model B)

When developing the sleep model for pretraining, we used the architecture that was used for Model A. However, we made several unique modifications. We created 18 duplicates of each 30-second sleep segment and added Gaussian noise (µ=0, σ=0.7) to the original segment and each duplicate. After concatenating each segment, samples input to the model were of the same dimensions of those of the basic model (i.e., 3000 time points by 19 channels). Our next major modification was adding a channel dropout layer [18] with a dropout rate of 25% at the beginning of the model to force the model to learn representations from all input channels rather than overfitting on only one channel. We used a training approach similar to that of [19]. We used a group shuffle split cross validation with an 80-10-10 training-validation-test split. We trained the model for a maximum of 200 epochs with a batch size of 128, an Adam optimizer (learning rate = 0.0075) with an adaptive learning rate that decreased by an order of magnitude after every 5 epochs without an increase in validation accuracy, and early stopping after 20 epochs without an increase in validation accuracy.

#### MDD Pretrained Model with Frozen Convolutional Layers (Model C)

To examine the utility of the features learned by Model B, we used the Model B weights from the sleep classification fold with the highest weighted F1 score, removed the channel dropout layer at the start of the model, froze the convolutional layer weights, re-initialized the dense layer weights, and decreased the number of nodes for the final softmax layer from 5 to 2. After making these modifications to the pre-trained model, we used an identical training approach to that used in the training of Model A.

#### MDD Pretrained Model without Frozen Layers (Model D)

After examining whether Model B had useful representations, we determined whether further optimizing Model C by unfreezing all weights would improve performance. We initialized Model D with the Model C weights and otherwise used the same training procedure as in Model A.

### C. Explainability Analysis

We lastly examined how pretraining affected the representations learned by MDD Models A, C, and D. As such, we applied an adaptation of the EasyPEASI spectral explainability approach [20] that was first presented in [21] and a zero-out channel ablation approach [22], [23].

Our spectral explainability approach involved converting test samples to the frequency domain permuting values within a given canonical frequency band across samples, converting back to the time domain, and calculating the change in BACC following perturbation. We used the δ (0-4 Hz), θ (4-8 Hz), α (8-11 Hz), β (12-25 Hz), and γ (25-50 Hz) frequency bands.

Our channel ablation approach involved iteratively assigning values for each channel in test samples to values of zero and examining the change in model performance. While we would have preferred to use the line-noise ablation approach described in [23], the use of publicly available data with a notch filter at 50 Hz removed line-related noise.

## III. Results and Discussion

In this section, we describe and discuss our model performance results, explainability results, and limitations and future work for this study.

### A. Model Performance Comparison

Model B obtained modest levels of performance (Table 1) below that of existing sleep stage classification studies [19], [21]. Nevertheless, classification performance was above chance-level for most classes. This is important as it shows that our use of duplicate channels with additive noise and channel dropout did not keep the model from learning. Model A shows the baseline level of performance that could be expected from our architecture and data (Table 2). All metrics were above 75%, with SENS being significantly higher than SPEC and BACC being close to 85%. Interestingly, model C, which just used the features extracted by model B, was able to obtain modest levels of classification performance. Model B had comparable SENS and BACC and lower SPEC relative to model A. Importantly, our final model, Model D, demonstrated a significant improvement over baseline, having comparable SENS, greatly increased SPEC, and modestly increased BACC. These findings support the viability of our pre-training approach for improving model performance.

**Table 1.** Model B Performance Results.

|  | F1 | SENS | PREC |
| --- | --- | --- | --- |
| **Awake** | 87.43±05.85 | 78.94±08.79 | 98.76±01.06 |
| **NREM1** | 07.29±02.89 | 04.55±02.00 | 20.18±06.60 |
| **NREM2** | 71.90±06.03 | 87.06±06.10 | 61.47±06.63 |
| **NREM3** | 67.65±09.47 | 90.09±07.17 | 55.99±13.88 |
| **REM** | 28.72±16.36 | 23.79±16.33 | 40.12±16.66 |

**Table 2.** Performance Results for Models A, C, and D.

|  | BACC | SENS | SPEC |
| --- | --- | --- | --- |
| **Model A** | 84.19±11.21 | 92.50±18.46 | 75.88±22.81 |
| **Model C** | 80.33±16.96 | 93.00±14.92 | 67.67±33.03 |
| **Model D** | 87.59±13.27 | 91.69±17.46 | 83.49±18.54 |

### B. Explainability Analysis

As shown in Figure 2, all models greatly relied upon δ and γ activity and ignored θ and α activity. Interestingly, β was of greater median importance for Model C than Model A and for Model D than Model C. This would suggest that the model learned β activity from the sleep data that it was later able to capitalize on when applied to the MDD classification data. There were also some similarities in spatial importance, as shown in Figure 3. Namely, F7, Cz, and P4 were of importance across models. Nevertheless, there were some significant differences. In several instances, Model D became less like Model C and more like Model A (e.g., F8, Pz, Fz). However, in several instances Model D learned to rely on channels in a manner very distinct from both Models A and C (e.g., F4, O2).

**Figure 2.**
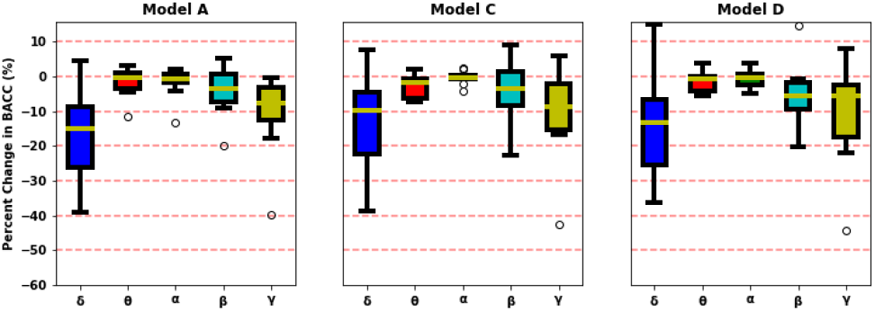
Spectral Importance. In panels from left to right are spectral importance for Models A, C, and D. The y-axes are aligned and show the percent change in BACC, and the x-axes indicate each frequency band. Note that δ, followed by γ activity was important across models, though the pretrained models also prioritized β activity.

**Figure 3.**
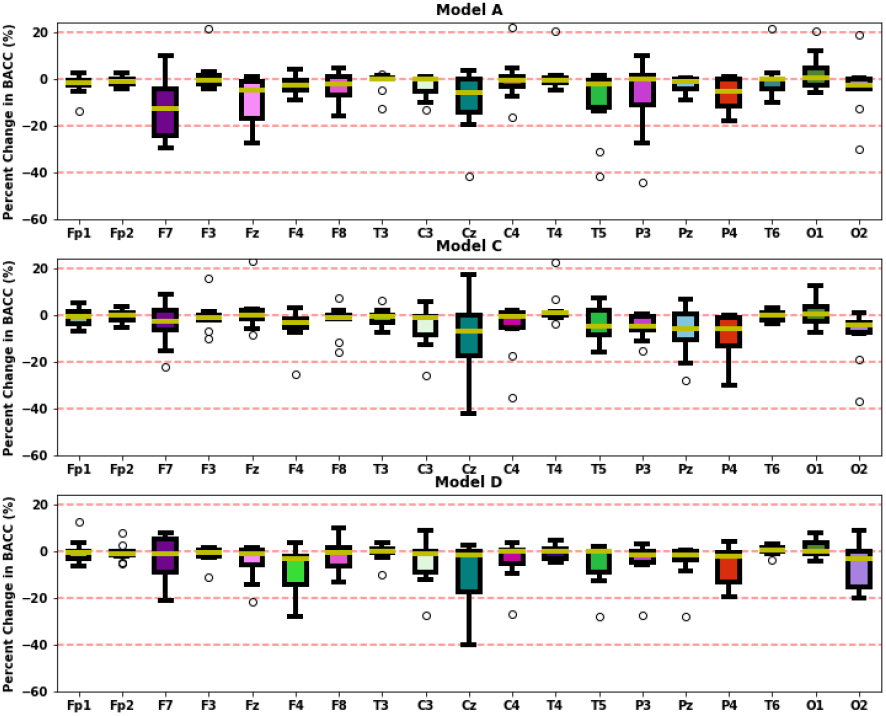
Spatial Importance. The top, middle, and bottom panels show spatial importance for Models A, C, and D, respectively. Channels are displayed on the x-axis, and the percent change in BACC is shown on the y-axis. Note how F7, Cz, and P4 are important across models. Model D was more like Model A than Model C in F8, Pz, and Fz importance, and Model D was unique in relying upon F4 and O2.

### C. Limitations and Next Steps

While our pretraining approach increased model performance and we examined the effect of pretraining upon learned representations, there are several avenues for further development and investigation of our proposed approach. In future studies, we will examine how the performance of our model upon the initial sleep stage classification task affects subsequent MDD model performance. We will further investigate how our pretraining approach interacts with our MDD data augmentation approach and investigate how each aspect of our proposed approach contributes to final MDD model performance. In each of these analyses, we will increase our number of folds above the traditional 10-folds to obtain statistically meaningful results.

## IV. Conclusion

Deep learning methods are increasingly being applied to raw EEG data. However, the relatively small size of many EEG datasets can make training reliable deep learning models a challenge. In this study, we propose a new raw EEG pretraining approach in which we transfer representations learned from single-channel sleep stage classification data to multichannel MDD diagnosis. Our proposed approach improves model performance, and we show through explainability analyses how pretraining affects the features learned by our models. Our proposed approach has the potential to expand the use of deep learning methods across more raw EEG datasets and lead to the development of more reliable EEG classifiers.

